# The Structural Basis of Genome Silencing: The Inactive X Chromosome Resolved by Electron Spectroscopic Imaging

**DOI:** 10.64898/2026.10.28.761900

**Authors:** Zahra Baghestani, Helena Alvarenga, Ren Li, Matthew Mcneil, Eden Fussner-Dupas, Mike Strauss

## Abstract

The eukaryotic nucleus has a hierarchical organization that permits genomic regulation. The genome is packaged as chromatin, which is silenced conditionally as facultative heterochromatin, or more stably silenced as constitutive heterochromatin. The smaller fractions of the genome that contain gene-rich sequences are organized into open chromatin conformations known as euchromatin. The inactive X (Xi) chromosome territory presents a uniquely powerful model system to investigate chromosome-scale gene silencing, as the entire chromosome undergoes coordinated epigenetic repression. The unique chromosome-wide silencing event provides a tractable system for investigating how molecular mechanisms of gene repression are translated into mesoscale structural features. This work builds on previous structural studies of the Xi that have leveraged conventional electron microscopy (EM) and cryo-EM, which have led to remarkable insights into the structural underpinnings of genome silencing. These approaches, however, are unable to directly distinguish chromatin from RNA and protein features. To resolve the molecular composition of the Xi chromosome territory and to provide a model of the structural organization of the chromatin, RNA and protein we analyzed the Xi using Electron Spectroscopic Imaging (ESI). We are able to resolve structural features without the use of contrast enhancing stains, using the distinct elemental signatures of chromatin, RNA and protein and combining them into composite images. By coupling ESI and electron tomography, we can circumvent the projection limitation of EM and provide an unprecedented view of the spatial organization of chromatin fibres in relation to structural features including RNA and proteins in 3D. Additionally, we employed correlative light-microscopy with ESI in physical serial sections and computationally assembled a structural model of the entire Xi chromosome territory revealing an intricate spatial network with distinct chromatin domains, distributed RNA-protein clusters, and identified a large, and previously uncharacterized mesoscale non-chromatin structure. Together, these observations provide new insights into the architecture of the Xi chromosome territory and shed electrons on the structural basis of chromosome-scale genome organization.

## Introduction

Genome regulation emerges from the hierarchical organization of chromatin, RNA, and protein assemblies within the eukaryotic nucleus. DNA is packaged as chromatin, where DNA polymers wrap around core octamers of histone proteins forming beads-on-a-string 10 nm chromatin fibres. Several non-histone proteins associate conditionally with chromatin and are integrally involved in genome regulation. The remaining nuclear space is occupied by diverse RNA- and protein-rich nuclear compartments. Through this coordinated organization, transcription is regulated, and genome stability is maintained.

This regulatory architecture can be conceptualized across multiple structural scales. At the molecular scale, organization arises from interactions between DNA, nucleosomes, chromatin-associated proteins, and epigenetic modifications. At the chromosome scale, entire chromosome territories exhibit cell-specific conserved spatial organization and contribute to genome regulation. Studies spanning structural biology, genomics, and imaging have established key principles governing genome organization, including nucleosome structure^1^, chromatin regulation^2^, chromosome territory formation^3^, chromatin looping^4^, topologically associating domains^5^, and active and inactive chromatin compartments^6–8^. Despite these advances, considerably less is known about the physical organization of chromatin at an intermediate, or mesoscale, where individual chromatin fibres and associated protein-RNA assemblies contribute to chromosome-scale nuclear architecture and genome regulation^9^. Here we use the term mesoscale to describe structural features spanning tens to hundreds of nanometers that bridge local chromatin organization and chromosome-scale architecture.

Understanding nuclear organization at the mesoscale is particularly important for determining how heterochromatin contributes to genome regulation. Heterochromatin domains can span millions of base pairs. These domains contribute to transcriptional repression, genome stability, and nuclear organization. Constitutive heterochromatin, which includes centromeric, telomeric, and repetitive genomic regions, remains relatively stably repressed across cell types, whereas facultative heterochromatin can be established to conditionally silence a genomic region and reversed dynamically during development or in response to environmental cues (reviewed in Allshire *et al.*^10^). Although many proteins and RNAs that are involved in heterochromatin formation and maintenance have been identified ^11–13^, considerably less is known about how these different components are assembled into mesoscale structures. In particular, the physical relationship between chromatin fibres and the surrounding protein- and RNA-rich environments characteristic of heterochromatin domains remains poorly understood.

Among heterochromatin structures, the inactive X chromosome (Xi) represents a uniquely powerful model for investigating chromosome-scale silencing. During X chromosome inactivation, one X chromosome is transcriptionally silenced through the establishment of a large facultative heterochromatin domain enriched in Xist, Polycomb-associated chromatin modifications, and numerous silencing-associated proteins ^14–18^. Because an entire chromosome undergoes coordinated epigenetic repression while remaining structurally distinguishable within the nucleus, the Xi provides a tractable system for investigating how molecular mechanisms of gene repression are translated into mesoscale structures.

Over the past two decades, studies of the Xi have fundamentally advanced our understanding of chromosome architecture and gene silencing. Chromosome conformation capture approaches (HiC)^19^, super-resolution imaging^20^, conventional transmission electron microscopy^21^, and genomic analyses have revealed that the Xi is a highly structured chromosome territory, rather than a uniformly compacted chromatin mass. These studies characterized the Xi as having two megadomains separated by the DXZ4 locus with long-range superloops involving DXZ4, FIRRE, and other architectural loci, compartmentalized chromatin environments, and regions that escape transcriptional silencing ^19,22,23^. Furthermore, distinct chromatin states, including H3K27me3-rich facultative heterochromatin and H3K9me3-enriched constitutive heterochromatin associated with repetitive DNA elements, occupy different regions of the chromosome territory ^24^. Taken together, these observations suggest that mesoscale chromatin folding contributes to Xi chromosome organization and provides an important link between chromosome architecture and gene regulation^25,26^.

Despite these advances, current models of Xi chromosome organization are predominantly derived from chromosomal interaction frequencies or the localization of specific proteins and RNA molecules. These approaches have transformed our understanding of chromosome topology, yet they provide limited information regarding the physical composition and arrangement of the chromosome territory itself. Chromosomal interaction maps identify genomic loci that are associated with one another but do not directly reveal how chromatin fibres, RNA, and protein-rich assemblies are physically arranged within the chromosome. Likewise, genomic perturbation studies and fluorescence imaging approaches require candidate molecules to be identified a priori. Consequently, our understanding of the physical organization of chromatin and non-chromatin structures within the Xi, remains incomplete. Recent advances in cryo-electron tomography (cryoET) have begun to bridge this gap by visualizing near-native chromatin within the inactive X chromosome at nanometer scales. These studies demonstrate that Xi organization is built on progressive chromatin compaction across multiple structural scales from individual nucleosomes to larger chromatin domains, rather than through the formation of canonical higher-order chromatin fibres (reviewed in ^27^). These observations are consistent with previous studies of mitotic chromatin analyzes by small-angle x-ray scattering, and electron spectroscopic imaging of mouse tissues ^28,29^. Chromatin within the inactive X chromosome is organized into discrete domains enriched in histone H1 chromatosomes ^30–32^, and hyperacetylation disrupts Xi chromatin domain boundaries. Developmental studies further demonstrate that dynamic structural transitions accompany the establishment of Xi chromosome inactivation and highlight critical roles for SPEN, histone deacetylation, and Xist-associated pathways in driving Xi architecture in iPSC models. Collectively, these studies provide unprecedented insights into local chromatin organization and the molecular mechanisms underpinning chromosome compaction.

When considered holistically, the models emerging from HiC and cryoET reveal an important and outstanding question. While Hi-C defines chromosome topology, and cryoET reveals chromatin architecture at near-molecular resolution, considerably less is known about how chromatin and non-chromatin structures are physically integrated across an entire chromosome territory. It remains unclear whether molecularly defined Xi domains correspond to physically distinct structural compartments, how chromatin fibres are organized within those compartments, and whether chromosome-scale organization arises solely through chromatin organization or if it involves larger macro-molecular assemblies of chromatin, protein and RNA. These questions are particularly relevant given mounting evidence that key Xi-associated proteins and RNA complexes are required for both the establishment and maintenance of chromosome silencing, and that several Xi-associated proteins exhibit condensate-like behaviour. Together, these observations raise the tantalizing possibility that large-scale Xi architecture may be scaffolded in part through mesoscale protein or protein-RNA assemblies. Addressing these questions requires an approach capable of simultaneously resolving chromatin organization, non-chromatin structures, and chromosome-scale architecture.

Herein, we use electron spectroscopic imaging ^33^ to investigate the structural organization and physical composition of the inactive X chromosome in low-passage female human fibroblasts. By directly distinguishing chromatin-rich, RNA-rich and protein-rich structures *in situ*, we identify a hierarchical organization of the inactive X chromosome. These analyses reveal structurally distinct chromatin compartments characterized by discretely organized 10 nm chromatin fibres within facultative heterochromatin domains. We further provide a chromosome-scale view of how chromatin and non-chromatin structures are integrated within the inactive X chromosome territory and identify a previously unrecognized mesoscale non-chromatin structure of 300 nanometers of diameter, spanning approximately 1.5 micrometers. For context on the scale of these structures, the entire nucleus of these cells is ∼23 micrometers. Jointly, these observations provide new insights into the architecture of the inactive X chromosome and the structural basis of chromosome-scale genome organization.

## Results

### Facultative chromatin within the inactive X is less dense than constitutive heterochromatin

To visualize the impact of large-scale gene inactivation that characterizes the inactive X chromosome, we measured both the chromatin fibre density and protein distribution in Wi-38 female fibroblasts using electron spectroscopic imaging (ESI). Human Wi-38 female fibroblasts were selected as our model system, as they are non-transformed, have stable genomes, and well-characterized and observable Barr bodies enriched in histone H3, tri-methylated lysine 27 (H3K27me3). This histone modification is a canonical emblem of facultative heterochromatin, which, in the case of the inactive X chromosome, is established, and maintained over cell generations. Wi-38 cells were embedded and thinly sectioned (70 nm) in preparation for electron microscopy. The cells were immuno-labeled with antibodies against H3K27me3, and imaged on-section, to identify the approximate boundaries of the large facultative heterochromatic region within the Xi by fluorescence microscopy. After subsequent transfer of the sections to an electron microscope, the Xi was imaged at several magnifications by ESI, first using a low magnification ESI phosphorus-enhanced image to align the fluorescence image to the high magnification images, and then at higher magnification for elemental mapping by jump-ratios. Two examples of Xi in two nuclei are presented in Figure 1, with the phosphorus signal in yellow, and the nitrogen signal in cyan (Fig 1A&B iv). Correlative LM-ESI is described in full in Bazett-Jones, et al, 2008 ^34^.

**Figure 1.**
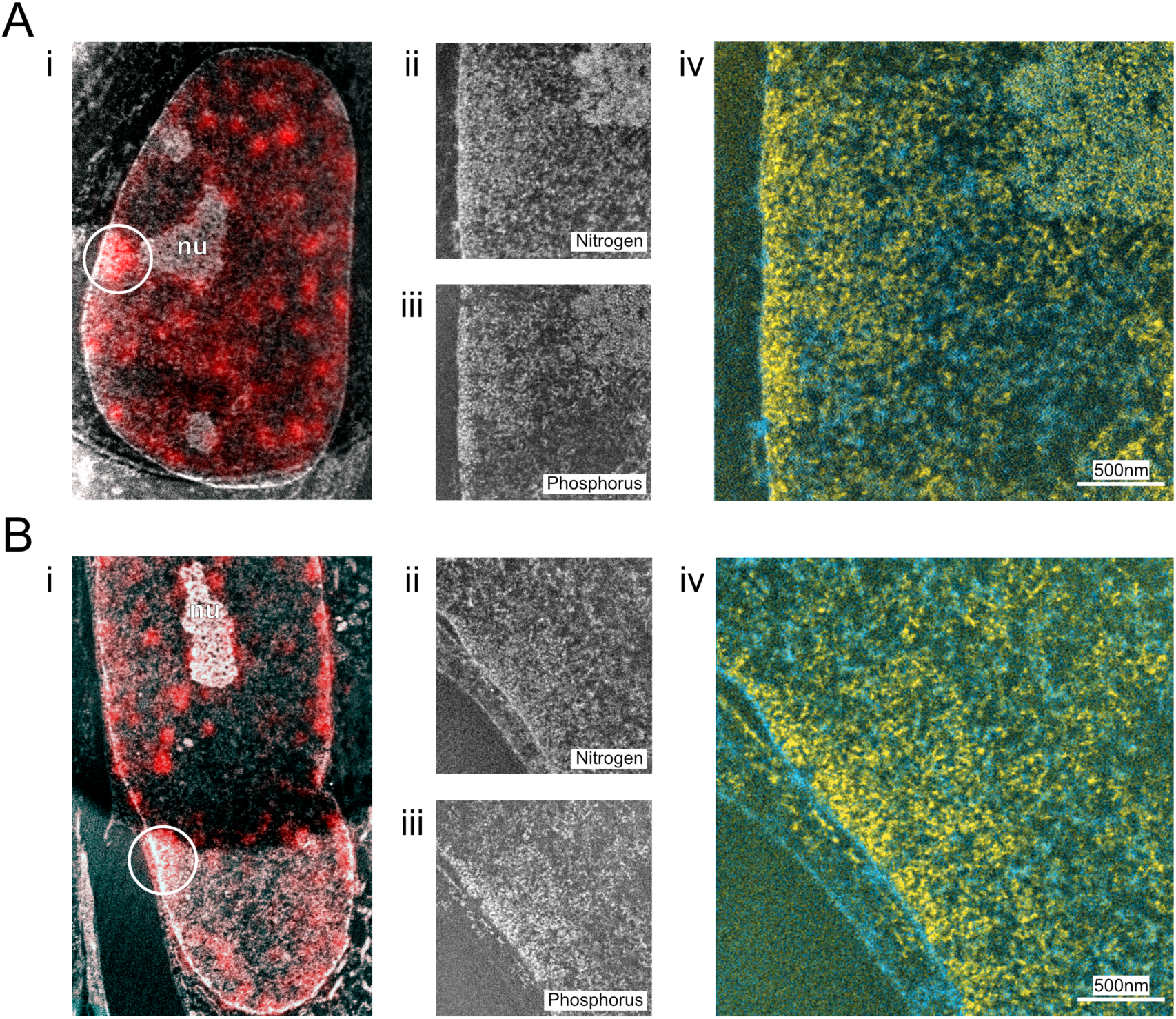
Representative images from two separate nuclei (panel **A** and **B**) showing results from correlative LM-ESI analysis of the inactive X chromosome in Wi-38 cells. i) On-section fluorescence imaging, acquired by epifluorescence, of Wi-38 to detect H3K27me3-enriched chromatin regions. The circle highlights the region containing the Xi chromosome territory imaged at higher magnification. ii) Two-window nitrogen mapping provides a quantitative image of nitrogen-rich structures within the nuclei. iii) Two-window phosphorus mapping of the same region as ii) reveals the phosphorus-containing structures in this region of the nucleus. iv) Phosphorus-subtracted nitrogen image in cyan is overlaid with the phosphorus image in yellow. Structures shown in cyan represent protein-rich regions which are concentrated in nitrogen above the background of chromatin (which is a protein-nucleic acid complex).

Phosphorus and nitrogen maps can be used to distinguish chromatin-based, RNP-based, and protein-based structures in the nucleus described in detail in ^33^. Briefly, as chromatin consists of very long polymers of nucleic acids which wrap around the protein-rich histones in strings of nucleosomes, by using phosphorus and nitrogen element mapping, chromatin is readily visualized as phosphorus-rich structures. In nucleic-acid poor or chromatin excluded regions, which is a hallmark of nuclear bodies, nuclear structures are enriched in nitrogen above the background of the nitrogen concentrations observed in chromatin. RNA is distinguished morphologically from chromatin, by assessment of the diameter and organization, and by phosphorus-to-nitrogen ratio analyses, as these two polymers vary significantly by both of these parameters. Chromatin associated within the H3K27me3-enriched regions of the Xi chromatin are more compact, compared to the dispersed open chromatin in nucleoplasmic space. The chromatin at the edge of the nuclear envelope is comprised of the constitutive heterochromatin that is largely made up of repetitive elements found throughout the human genome and in regions attached to the H3K27me3-enriched domain. We assume this to be a part of the Xi chromosome territory. Compact constitutive heterochromatin lines the nuclear envelope in most mammalian cells, and, outside of the Xi domain, is likely to be contributed by different chromosome territories. Using ESI, constitutive heterochromatin is readily visible and is interspersed with protein-rich nuclear pore complexes, which are resolved as nitrogen-dense baskets protruding from gaps in the nuclear peripheral constitutive heterochromatin into the cytoplasmic space. The gaps in the constitutive heterochromatin opposite the nuclear baskets often extend several hundred nanometers into the nucleus. Constitutive heterochromatin domains appear more phosphorus dense than the facultative regions of the Xi chromosome territory.

To quantify the chromatin density associated with the Xi, we measured the phosphorus density by ESI from the H3K27me3-enriched facultative heterochromatin, the highly compact constitutive heterochromatin chromatin at the nuclear periphery, and the dispersed open euchromatin within the nucleoplasmic space from images taken of 91 sections from 7 different cell nuclei (Figure 2). The phosphorus density in the facultative, constitutive and euchromatin regions was significantly different, with the constitutive heterochromatin demonstrating the highest phosphorus density (6.39P/nm^2^), and the open euchromatin with the least phosphorus density (4.45 P/nm^2^). Facultative heterochromatin of the Xi chromosome territory had an intermediate phosphorus density (5.17 P/nm^2^) with average compaction values being approximately 80% of that measured in constitutive heterochromatin domains.

**Figure 2.**
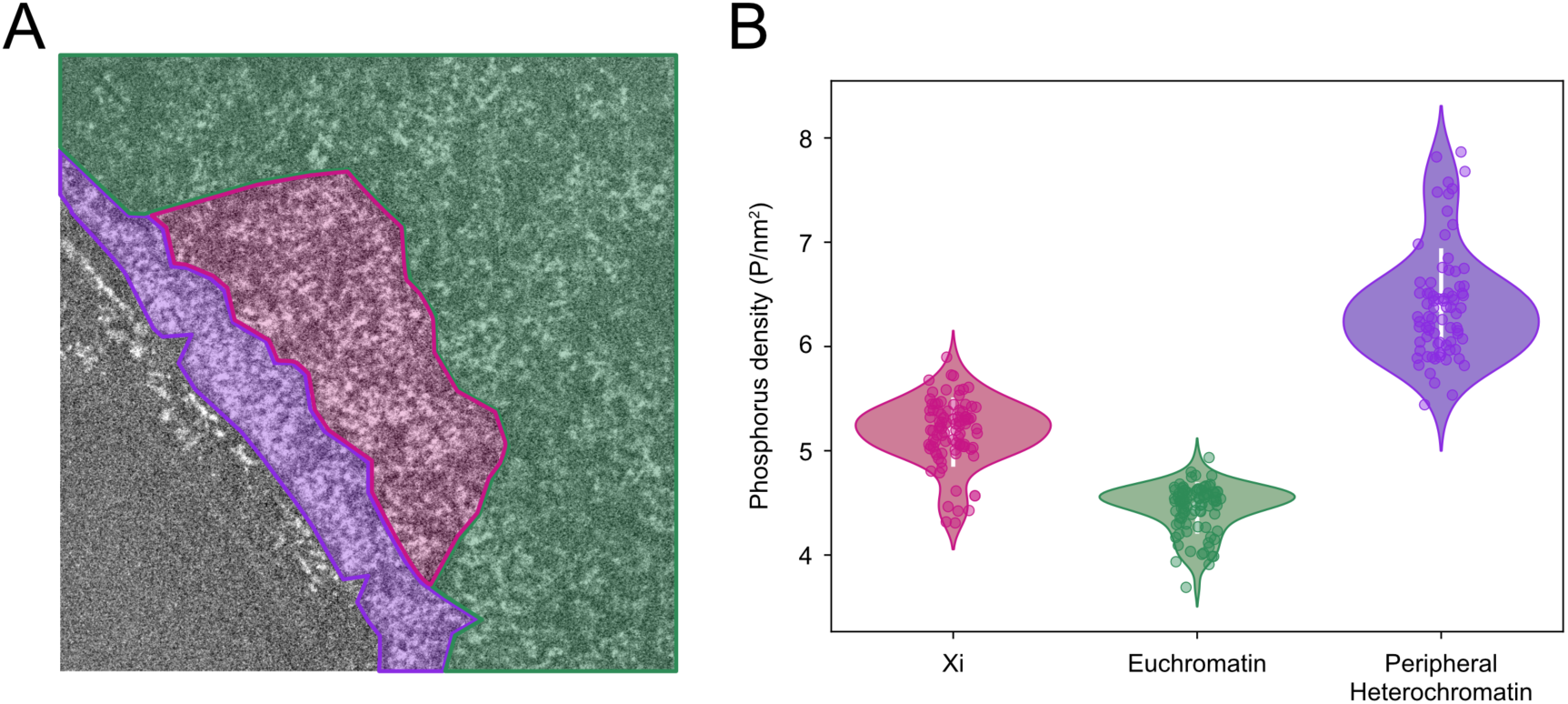
A segmented example of a region within a Wi-38 cell nucleus with three distinct chromatin sub-types (**A**) and aggregated phosphorus density analyses from 91 physical sections. The phosphorus density (**B**) in the dispersed euchromatin and that of the compact constitutive heterochromatin domains was significantly different from that found in the Xi (p values of 7.74 x 10^-26^ and 1.68 x 10^-28^, respectively). Constitutive heterochromatin regions have the most compaction, with mean chromatin density values of 6.39 phosphorus per nm^2^. The mean Xi chromatin density is 5.17 P/nm^2^, about 80% of the density of constitutive heterochromatin. The chromatin density of Xi is higher than that of the dispersed euchromatin in these cells at 4.45 P/nm^2^.

The phosphorus density analyses were performed on thin, 70 nm sections. Since transmission electron microscopy compresses all the data into a 2-dimensional image plane, and both chromatin and RNA contain one phosphate atom per nucleotide, it remains possible that the density values calculated above contain both chromatin and RNA-based structures. Especially in light of the known contribution of RNA in directing and potentially maintaining the Xi in the active state, our density measures could be inflated through RNA phosphorus contributions. By collecting 3-dimensional data chromatin fibres can be measured independently from RNA polymers, therefore, to circumvent the projection limitation which compresses these data into a single-image plane.

### 3-Dimentional (3D) analysis of the Xi chromosome territory is an integrated network of chromatin and large protein assemblies

To visualize the potential protein components in 3D, and distinguish chromatin from RNA within the Xi chromosome territory, we utilized ESI tomography, which simultaneously enables accurate chromatin fibre measurements.

We acquired four energy-loss images (at 120eV, 155eV for P, and 385eV, 415eV for N) at each tilt over a 110-degree tomographic tilt range in 2-degree increments and recombined these data using IMOD. The processing procedure and required linux shell commands to assemble two-element 3D ESI-tomographic reconstructions is described in Fussner *et al* ^34^. This approach enabled us to measure chromatin fibres, protein, and RNA molecular assemblies within the Xi simultaneously. In these analyses (Figure 3), we observe that the chromatin fibres themselves are organized in 10-nm chromatin configurations, consistent with our previous observations of chromatin in mouse tissue culture cells and in mouse tissues ^28^. RNA polymers are distributed throughout the chromosome territory but are not more visibly concentrated within the facultative region of the Xi relative to the surrounding chromatin regions. An unexpected observation which emerged from these reconstructions was the presence of protein-rich fibre-like structures both within and surrounding the Xi chromosome territory, which were noticeable between chromatin fibres, sometimes in regions where chromatin is clearly excluded in the physical section. These protein-rich structures had varying diameters and persistent lengths.

**Figure 3.**
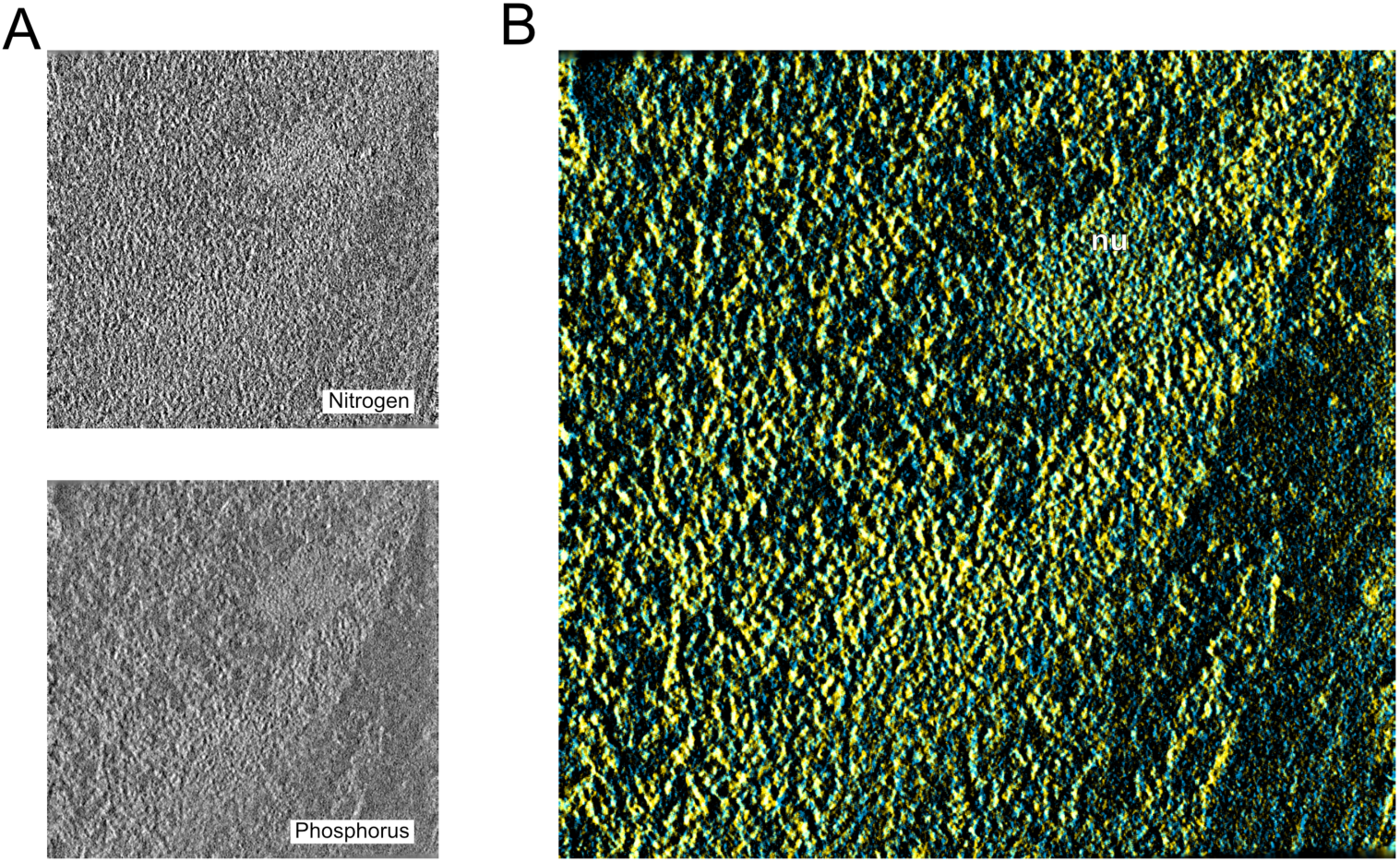
Electron spectroscopic tomograms separate elemental signals (A: nitrogen (top), phosphorus (bottom)) along the Z-direction. The overlay (phosphorus – yellow; nitrogen – cyan) of 5nm slabs through the tomogram show the spatial and structural relationship between chromatin and nitrogen-rich nuclear structures. The Xi chromatin territory is at the nuclear envelope and is associated with the nucleolus (nu). Structures that are clearly visible in cyan have more nitrogen than the background signal associated with chromatin fibres themselves.

Dual element tomography provided unprecedented structural insights into chromatin structure and organization, yet the nature of these protein-rich and chromatin depleted structures remains only partially resolved, as the tomographic reconstructions are done on thin, 70 nm structures, and these protein-rich structures may extend beyond this scale. To address this possibility, we assembled an entire 3D view of Xi from 2D ESI maps of sequential serial sections. These serial sections were first imaged for the Xi using H3K27me3 fluorescence, and low and high magnification ESI images were acquired of each physical section. These data were computationally recombined using a combination of low magnification overviews to guide positioning and Midas ^35^ (for aligning serial sections). The Xi and protein feature were segmented by hand in IMOD.

This approach provided a complete sampling of the various structural and domain-based features of the Xi chromosome organization. H3K27me3 is a canonical epigenetic histone modification associated with facultative heterochromatin, which means several regions within a given 70 nm slice of the nucleus are antigenic and have clustered fluorescence signal. However, due to the size and scale of the Xi, the Barr bodies are readily identified by mapping continual fluorescence signals within the same region of interest over multiple sections.

This methodology enables us to capture the emerging edge of the chromosome territory, where a relatively small amount of H3K27me3 is present in the section at the position of the Barr body. The domain containing the H3K27me3 s gnal is barely recognizable as a distinct structure in the corresponding ESI micrograph. As sections begin to capture structures in the middle of the Xi territory, more of the volume is dominated by H3K27me3-enriched chromatin. Consistent with the findings of Chadwick and Willard ^36^, the Xi chromosome territory is located immediately adjacent to the nuclear envelope (5 of 7 full reconstructions) or adjacent to the nucleolus. The chromatin enriched in H3K27me3 can be distinguished both from the surrounding euchromatin in the nucleoplasmic background and from the constitutive heterochromatin at the nuclear envelope, by virtue of the distinctive phosphorus density of facultative heterochromatin. We expect that in some sections, the constitutive heterochromatin at the nuclear envelope is derived from the Xi itself, while remaining physically distinct, yet still attached, from the H3K27me3-rich component of the Xi.

The combined 3D model of the entire Xi territory reveals some surprising structural features. Not only is the chromatin fibre density non-uniform due to the segmentation of constitutive, facultative and euchromatin within the whole territory; we observe frequent chromatin-free holes and channels. An example of these protein-rich architectural features is shown in Figure 4A,B. We observed several distinct and diverse non-chromatin structures within the Xi chromosome territory, including RNPs, which are both morphologically distinct and have a much higher nitrogen to phosphorus ratio than chromatin, both within and surrounding the Xi periphery (Figure 4B). The entire Xi territory contains three different types of chromatin measured by density, and regions of concentrated protein-rich fibrous structures. The most unexpected observation was the presence of a huge protein-rich structure with a 300 nm diameter spanning nearly the entire length of the chromosome territory, which is at least 1.5 microns in length. It is worth noting that in this example the Xi territory is not attached to either the nuclear envelope or the nucleolus, and the chromatin from this territory appears to be anchored around this protein-rich structure. In our models of two other Xi territories which are associated with the nuclear envelope we observed protein-rich structures of this same diameter, however, in these inactive X chromosome territories the protein-rich structures were located towards the edge of the chromosome territory, rather than through the centre core as observed in this example. Electron spectroscopic imaging, combined with three-dimensional resolution approaches, have provided a means to resolve, in a quantifiable way, the contributions of chromatin, RNA and protein structures within and surrounding the Xi chromosome territory.

**Figure 4.**
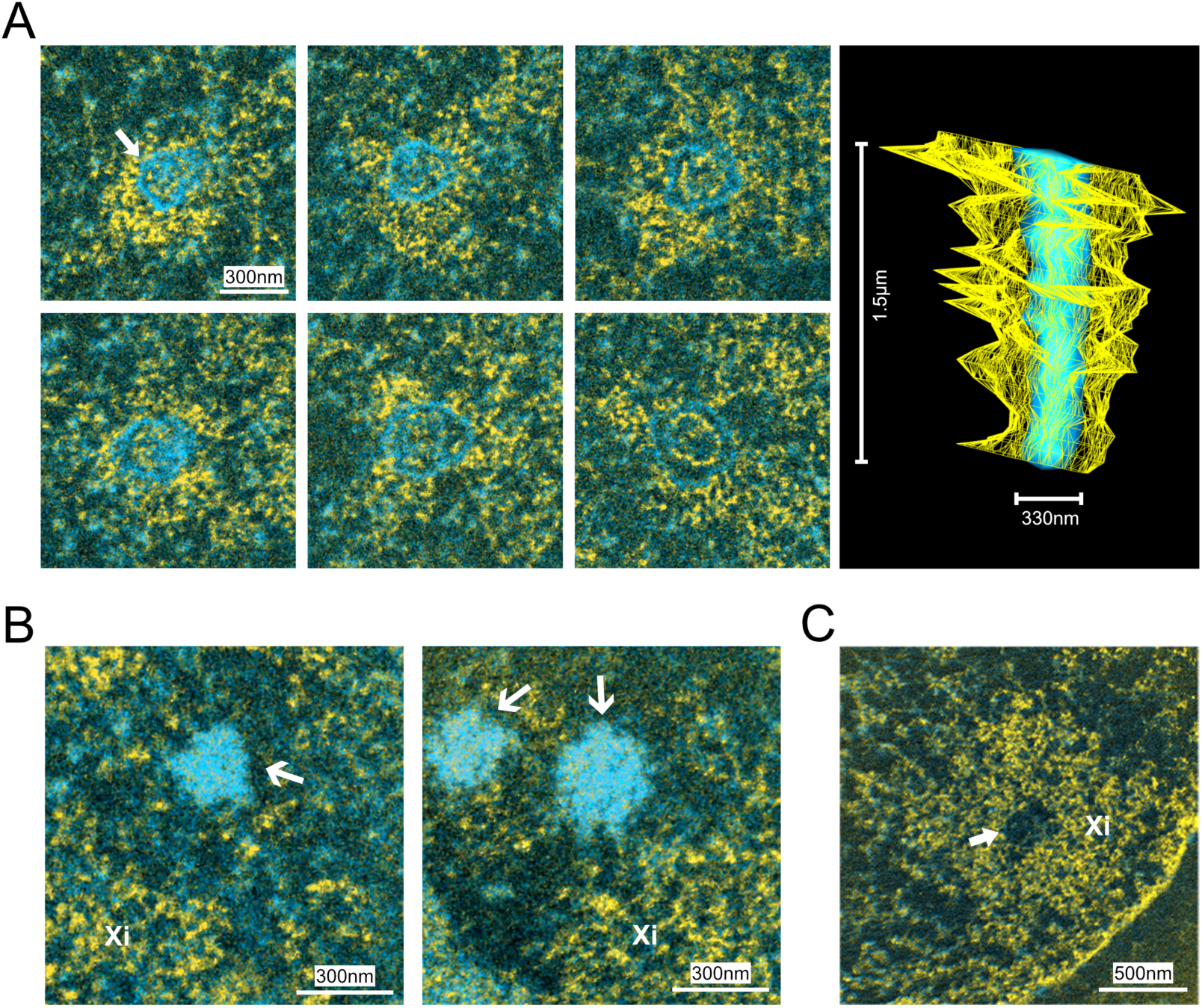
Large scaffolding non-chromatin structures are observed within and proximal to the Xi chromosome territory. Shown in panel **(A)**, **(B)** and **(C)** are four different Xi chromosome territories that have been modeled from physical serial section ESI image processing. The nitrogen signal has been subtracted from the phosphorus image in each physical section, structures that appear cyan in these composite images are more concentrated in protein than the bulk chromatin. Arrows in panels **(A)** and **(B)** indicate the very large protein-rich structures that are observed in the core of the Xi chromosome territory, while arrows in panel **(C)** show a phosphorus-poor pocket in the centre of the Xi territory. The model (top right) shows the contours that bound both the nitrogen-rich protein core and the H3K27me3-enriched chromatin mapped by correlative LM-ESI imaging.

## Discussion

Taken together, our results using Electron Spectroscopic Imaging (ESI) of the inactive X chromosome from Wi-38 human low passage fibroblasts, uncover a new layer of structural architecture associated with this chromosome territory. Using correlative ESI, ESI-tomography, and a 3D modeling approach, we were able to resolve structural features at several scales of organization. We find that the chromatin fibres within this cell type are comprised of 10 nm chromatin fibres, which are arranged with different degrees of compaction. Chromatin associated with the large facultative heterochromatin region within the Xi chromosome territory adopts an intermediate density state relative to constitutive heterochromatin and open euchromatin. Although we cannot rule out the possibility that the intermediate density measured here is a cell-type dependent phenomena, if more broadly reflective of Xi territories in other cell types, this suggests that efficient genome silencing does not require the degree of compaction observed in constitutively silenced genomic regions.

The inactive X chromosome is an integrated assembly of these three chromatin architectures and involves three potentially interesting macromolecular assemblies that are chromatin associated, but independent. The first, unsurprisingly, is RNA which in the cell rapidly associates with RNA-binding proteins to form small RNA clusters or domains (RNPs). RNPs are observed both embedded throughout the Xi territory and at its periphery. This distribution is consistent with super-resolution imaging studies of Xist, the long non-coding RNA responsible for initiating chromosome X inactivation, which revealed that Xist is organized into discrete foci distributed throughout the inactive X territory rather than forming a uniform coating of the RNA at the surface of the chromosome territory. Although Xist is indispensable for establishing the inactivation of Xi, its requirement for maintenance of silencing appears to be context dependent. In cycling human fibroblasts, the presence of RNA-containing RNPs throughout the Xi territory is consistent with ongoing Xist association, however, the present study did not directly assess whether or not the observed RNPs is Xist, and if this association is required for the maintenance of chromosome-wide gene silencing in this cell type ^20,37^.

We generated 3D models of the Xi, both using combined phosphorus and nitrogen ESI-Tomography, and by modeling and combining 2D data from serial sections of embedded cells through the entire Xi chromosome territory. These data demonstrate some unexpected architectural structural features of this large chromosome, which is enriched in conditionally silenced facultative heterochromatin. The presence of several protein-rich structures was surprising, as some spanned the entire length of the chromosome territory. The enriched nitrogen-signal and, in several instances, exclusion of the chromatin from these meso-scale protein assemblies, coupled with the fact that these cells are still cycling, leads us towards speculating that this sizeable feature (larger than a small nucleolus in these same cells) may serve as a scaffold for the chromatin in the H3K27me3 enriched regions to help stably promote gene silencing. The sheer scale of these structures may also explain why regions of the Xi chromosome are highly discrete and separated domains when examined by conformational capture approaches. Presumably, crosslinking between genomic regions separated by these nitrogen-rich protein structures would be unfeasible. It is tantalizing to speculate that the organization of the inactive X chromosome territory may in fact be, at least partially, driven by a large protein core that physically separates chromatin domains under these conditions. Cylcling cells must reorganize the nucleus completely during cell division, and reestablish the architectures necessary for chromosome territory organisation. The features we see here are thus unlikely to be the result of protein aggregation or uncontrolled maturation of a phase separated component. This conclusion may also provide some clues as to what biophysical phenomena is underlying the formation of these structures. Nuclear bodies, such as promyelocytic leukemia nuclear bodies (PML bodies) or the nucleolus, are approximately 3-times more nitrogen-rich than the rest of the nucleoplasmic space, and their formation is thought to be driven by phase separation due to the heightened protein concentration that occurs within non-membrane bound protein-rich sub-compartments. Many of these smaller channels and the meso-scale protein-rich scaffolding structures rival the protein concentrations observed in these other well-characterized nuclear bodies.

## Materials and methods

### Cell culture and sample preparation

Wi-38 human fibroblasts from the American Type Culture Collection (ATCC) were maintained in Dulbecco’s Modified Eagle Medium (DMEM) (Wisent, Québec, Canada) supplemented with 10% Fetal Bovine Serum (FBS) (FBS, Hyclone, Logan, UT, USA), 2mM L-glutamine, 10 μg/ml penicillin-streptomycin antibiotics at 37°C in 5% CO2. Cells were cultured on sterilized glass coverslips in 6-well plates. Confluent Wi-38 cells were maintained 10 days after plating to obtain the greatest percentage of quiescent cells with a distinct Barr body [4].

### Immunofluorescence labeling

Cells were removed from the media, rinsed rapidly in PBS (Wisent, Québec, Canada), then fixed in 2% PFA (Electron Microscopy Sciences, Hatfield, PA, USA) in PBS for 20 minutes at room temperature. All subsequent steps were performed at room temperature. After fixation, cells were washed three times in PBS, permeabilised with 0.5% Triton X-100 (Bishop #9002-93-1) in PBS for 5 minutes, then washed three times in PBS. Cells were incubated with primary antibody in PBS containing 5% donkey serum (Jackson ImmunoResearch, West Grove, PA, USA) for 1 hour. Primary antibodies used were: rabbit anti-H3K27me3 rabbit anti-H3K9me3 (at 1:300 dilution) (cat. 07-499 and 07-442, respectively; Millipore, MA, USA). Samples were washed three times in PBS and incubated in 1:200 Cy3-labeled donkey anti-rabbit secondary antibodies (Jackson ImmunoResearch) for 1 hour. Cells were washed and mounted in antifade (Sigma-Aldrich Co. Cat No. P3130) containing 1mg/ml DAPI (Sigma-Aldrich Co. Cat No. D9542).

### Immunolabeling, silver enhancement, dehydration and embedding

All steps were performed at room temperature unless otherwise indicated. Cells were fixed in 2% PFA (Electron Microscopy Sciences,Hatfield, PA, USA) in PBS for 15 minutes, washed with PBS, permeabilized with 0.5% Triton-X 100 in PBS for 5 minutes, washed three times in PBS, incubated in primary antibodies (rabbit anti-H3K27me3, cat. 07-499.; Millipore, MA, USA) for 1 hour, and washed three times. Ultra-small immunogold secondary antibodies (donkey anti-rabbit Aurion Ultra small Immunogold, Electron Microscopy Sciences) were diluted (1:50) in PBS containing 5% donkey serum (Jackson ImmunoResearch, West Grove, PA, USA), and incubated for 1 hour. Blocking buffer was added for at least one hour. Secondary Cy3-labeled donkey anti-rabbit secondary antibodies (1:50) (donkey anti-rabbit Cy3, Jackson ImmunoResearch) were diluted in PBS containing 5% donkey serum and added to the coverslips incubating in gold-conjugated antibodies on coverslips, and incubated for 1 hour. Antibodies were washed with PBS and the cells were post fixed in 1% glutaraldehyde (Electron Microscopy Sciences, Hatfield, PA, USA) diluted in PBS for 5 minutes, followed by washes in PBS and then distilled water. To visualize the nanogold particles, samples were subject to a silver enhancement protocol according to the manufacturer’s instructions (Aurion R-Gent SE-EM, cat. 25521; Electron Microscopy Sciences) and developed for 40 min at 37°C. To make silver enhancement mix, 1 μl of initiator and 40 μl of activator were mixed (mix 1) and then 10 μl of mix 1 was added to 190 μl of developer, and left in room temperature for 10 minutes before placed it onto samples. Cells were washed in distilled water and dehydrated in a graded ethanol series (30, 50, 70, 90, and 100%) for 30 minutes at each step. Cells were placed in 100% ethanol at 4°C overnight, then washed three times for 30 minutes in 100% ethanol, and embedded in Quetol 651 resin (cat. 20440; Electron Microscopy Sciences) for 1 hour. During this time, quetol mix was prepared such that 35mL of quetol 651, 54ml of NSA, 11ml of NMA were stirred for at least two hours. (NSA Kit for quetol mix, cat. 14640; Electron Microscopy Sciences). 100 μl DMP-30 was added to 5ml of the quetol mix, mixed for half an hour. Quetol was discarded and replaced with 5ml quetol mix for each sample and incubated with shaking for 2 hours. This was repeated twice and incubated at 65-70°C for at least 24 hours.

### Sectioning and carbon coating

Blocks of the embedded samples were cut using a sharp razor on a hot plate with the temperature set to approximately 90°C and glued to support bullets. Blocks were cut into 70nm sections with an Ultracut UCT microtome (Leica Microsystems Inc).

### Correlative immunofluorescence microscopy and electron spectroscopic imaging

Immunofluorescence images of the sectioned samples were acquired with a Leica DMRA2 microscope with a CCD camera (Hamamatsu) and OpenLab 3.5.1. A 63X objective lens was used to identify the heterochromatin domains enriched in the mark of interest. Sections were coated with a thin layer of 3 nm carbon film to protect them from the electron beam. Electron spectroscopic images were collected from these carbon-coated sections.

### Electron spectroscopic imaging

Energy filtered electron micrographs were taken with a transmission electron microscope (Tecnai 20, FEI) fitted with a GATAN post column imaging filter operated at 200 kV. All pre and post images were recorded on a Gatan CCD camera or K2 CMOS sensor, and images collected using Digital Micrograph software (Gatan). Low magnifications images of the entire cells were obtained by applying a 155 eV filter. For phosphorus images, two pre-edge reference images were collected at 120 eV and a post-edge phosphorus enhanced image collected at 155 eV energy loss were recorded prior to and following phosphorus ionization, respectively. Pre-image was subtracted from post-image to obtain the net phosphorus image. The same method was used to create net nitrogen images, with pre and post-images taken at 385 eV and 415 eV, respectively.

### Image analysis

Image processing and analysis were performed using the Digital Micrograph (Gatan), ImageJ, and Adobe Photoshop. Phosphorus maps were subtracted from the corresponding nitrogen maps to normalize chromatin-based structures to zero. Phosphorus maps were overlayed onto the subtracted images generating the merged phosphorus and nitrogen images. In the merged images, phosphorus is pseudo-coloured yellow, and phosphorus-subtracted nitrogen is cyan. Therefore, in the final images yellow and cyan represent chromatin- and non-chromatin or protein-based structures.

### Phosphorus density analyses

To quantify chromatin compaction, phosphorus density analyses of unprocessed phosphorus maps of 91 cell sections were performed in Jupyter Notebook 7.2.2 with Python 3.10.12. Regions were masked with Imod 5.0.1, according to enrichment of H3K27me3 in correlative images. The phosphorus maps were normalized by the average phosphorus content of masked ribosomes. The statistical analysis of average phosphorus density per region was performed pairwise Xi-Euchromatin and Xi-Heterochromatin by Mann-Whitney U Test.

### Three-dimensional reconstruction of whole chromosome

The whole chromosomes were aligned from serial sections using Midas, a program within the IMOD package^35^. Since high magnification serial section images of Xi have little consistent information between them for correlation, low magnification images for each serial section were first aligned to one another, since they contained large portions of the nucleus that vary more slowly across sections. Subsequently, high magnification phosphorus-nitrogen overlays were correlated with the aligned low magnification image stack. The relative image transforms were stored and applied to the high magnification images to obtain aligned overlays at high magnification.

### Tomographic reconstruction

The detailed description of ESI tomography is available in Fussner-Dupas *et al* ^34^, including links to useful scripts, which can also be found at https://doi.org/10.5281/zenodo.10928384. Briefly, tilt series for pre- and post-edge images for each element were generated using newstack (from the IMOD package). For each element, the pre- and post-edge images stacks were treated as dual tilt image stacks for ease of processing in etomo. After rough alignment by cross-correlation for each tilt series, one tilt series was selected, typically the post-edge phosphorus stack, to complete the full tomographic alignment. The alignment parameters for this tilt series was copied to the other tilt series, with an extra alignment step just prior to final aligned stack generation, run outside etomo to account for potential drift between acquisitions. The jump-ratio aligned tilt series was generated using newstack, and used for SIRT reconstruction, typically with 10 or 15 iterations.

## Acknowledgements

The authors would like to express their deepest gratitude to Dr. David Bazett-Jones, whose thoughtful manner and generous spirit resonate to this day; this work is a tribute to his legacy. EFD, MS, and HA were supported by FRQNT-NOVA grant: FRQ-NT 2023-NOVA-327333, and MS and HA were supported by NSERC Discovery grant: RGPIN-2020-04837.

